# ATP Binding Facilitates Target Search of SWR1 Chromatin Remodeler by Promoting One-Dimensional Diffusion on DNA

**DOI:** 10.1101/2022.01.24.477290

**Authors:** Claudia C. Carcamo, Matthew F. Poyton, Anand Ranjan, Giho Park, Robert K Louder, Thuc Dzu, Carl Wu, Taekjip Ha

## Abstract

One-dimensional (1D) target search is a well characterized phenomenon for many DNA binding proteins but is poorly understood for chromatin remodelers. Herein, we characterize the 1D scanning properties of SWR1, a yeast chromatin remodeler that performs histone exchange on +1 nucleosomes adjacent to a nucleosome depleted region (NDR) at promoters. We demonstrate that SWR1 has a kinetic binding preference for DNA of NDR length as opposed to gene-body linker length DNA. Using single and dual color single particle tracking on DNA stretched with optical tweezers, we directly observe SWR1 diffusion on DNA. We found that various factors impact SWR1 scanning, including ATP which promotes diffusion through nucleotide binding rather than ATP hydrolysis. A DNA binding subunit, Swc2, plays an important role in the overall diffusive behavior of the complex, as the subunit in isolation retains similar, although faster, scanning properties as the whole remodeler. ATP-bound SWR1 slides until it encounters a protein roadblock, of which we tested dCas9 and nucleosomes. The median diffusion coefficient, 0.024 μm^2^/sec, in the regime of helical sliding, would mediate rapid encounter of NDR-flanking nucleosomes at length scales found in cells.

## Introduction

Eukaryotic genomes are packaged into chromatin, the base unit of which is the nucleosome. Both the position of nucleosomes on the genome and their histone composition are actively regulated by chromatin remodeling enzymes (Yen et al., 2012). These chromatin remodelers maintain and modify chromatin architecture which regulates transcription, replication, and DNA repair (Tessarz and Kouzarides, 2014). A particularly well-defined area of chromatin architecture is found at gene promoters in eukaryotes: a nucleosome depleted region (NDR) of about 140 bp in length is flanked by two well-positioned nucleosomes, one of which, the +1 nucleosome, sits on the transcription start site (TSS) (Bernstein et al., 2004; Lee et al., 2007; Xu et al., 2009; Yuan, 2005) and the nucleosome on the opposite side of the NDR, upstream of the TSS, is known as the -1 nucleosome. The +1 nucleosome is enriched for the non-canonical histone variant H2A.Z (Albert et al., 2007; Raisner et al., 2005). In yeast, H2A.Z is deposited into the +1 nucleosome by SWR1 (Swi2/Snf2-related ATPase Complex), a chromatin remodeler in the INO80 family of remodelers (Ranjan et al., 2013). The insertion of H2A.Z into the +1 nucleosome is highly conserved and plays an important role in regulating transcription (Giaimo et al., 2019; Rudnizky et al., 2016).

While the biochemistry of histone exchange has been characterized, the target search mechanism SWR1 uses to preferentially exchange H2A.Z into the +1 nucleosome is not yet understood. The affinity of SWR1 for nucleosomes is enhanced by both long linker DNA (Ranjan *et al*., 2013; Yen et al., 2013) and histone acetylation (Zhang et al., 2005), and both factors play a role in the recruitment of SWR1 to promoters. A recent single molecule study further showed that SWR1 likely exploits preferential interactions with long-linker length DNA by demonstrating that H2A.Z is predominantly deposited on the long-linker distal face of the nucleosome (Poyton et al., 2021), similar to what is observed *in vivo* (Rhee et al., 2014). It is possible that SWR1 first binds long-linker DNA and then finds its target, the +1 nucleosome, using facilitated diffusion (**Figure 1A**), as was previously suggested (Ranjan *et al*., 2013). In a hypothetical facilitated search process SWR1 would first find the NDR through a three-dimensional target search. Once bound, it is possible the entire SWR1 complex diffuses one-dimensionally on the NDR, where it can encounter both the -1 and +1 nucleosomes. Facilitated diffusion has been shown to be essential for expediting the rate at which transcription factors and other DNA binding proteins can bind their target compared to a 3D search alone (Berg et al., 1981; Elf et al., 2007; Hannon et al., 1986; Ricchetti et al., 1988; Von Hippel and Berg, 1989). Furthermore, recently published *in vivo* single particle tracking found that chromatin remodelers have bound-state diffusion coefficients that are larger than that of bound H2B, hinting at the possibility that they may scan chromatin, but those studies could not distinguish between remodeler scanning and locally enhanced chromatin mobility (Kim et al., 2021; Ranjan et al., 2020). It is not known, however, if SWR1 or any other chromatin remodeler can linearly diffuse on DNA, and therefore make use of facilitated diffusion to expedite its target search process. Additionally, SWR1’s core ATPase, like other chromatin remodelers, is a superfamily II (SF2) double stranded DNA translocase (Nodelman and Bowman, 2021; Yan and Chen, 2020); while there is no evidence for SWR1 translocation on nucleosomal DNA, it remains possible that SWR1 may undergo directed, instead of diffusional, movements on a DNA duplex in the absence of a nucleosome substrate.

**Fig. 1.**
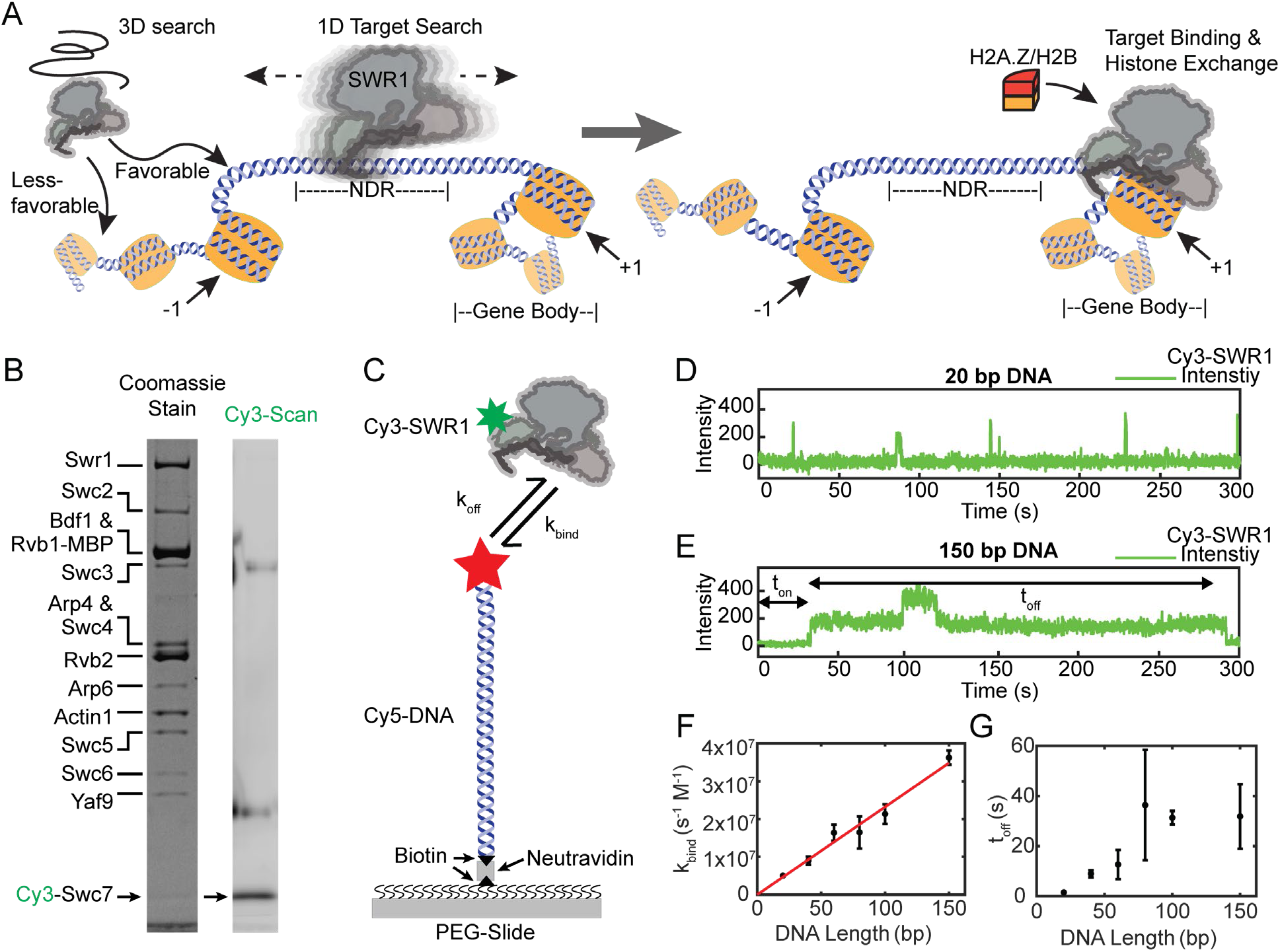
SWR1 binds DNA in short and long-lived states and prefers longer DNAs. (A) Proposed facilitated search mechanism for how SWR1 locates the +1 nucleosome. (B) A denaturing SDS-PAGE of reconstituted Cy3-SWR1 imaged for Coomassie (left) and Cy3 fluorescence (right). Cy3-Swc7 is faint when stained with Coomassie but is a prominent band in the Cy3 scan. The two diffuse bands that run at higher molecular weight and appear in the Cy3 scan are carry over from the ladder loaded in the adjacent lane. (C) A schematic for the single-molecule colocalization experiment where the kinetics of Cy3-SWR1 binding to Cy5-labeled DNA of different lengths was measured. (D-E) Representative trace for Cy3-SWR1 binding to (D) 20 bp Cy5-DNA, and to (E) 150 bp DNA. A second Cy3-SWR1 can be seen binding at approximately 100 s. (F) Measured binding time (k_bind_) for SWR1 to DNA of different lengths, error bars are standard deviation. The red line is a linear fit to the data, where R2 = 0.99. (n = 2 technical replicates) (G) The lifetime (t_off_) of Cy3-SWR1 bound to DNAs of different lengths, error bars are standard deviation. (n = 2 technical replicates) **Figure 1 – Source data 1** Numerical data underlying panel F and G **Figure 1 – Source data 2** Gel images (Coomassie and Cy3 scans) shown in panel B

In this study, we used a site-specifically labeled SWR1 complex to demonstrate that SWR1 can scan DNA in search of a target nucleosome. First, we characterized the kinetics of SWR1 binding to DNA and found that the on-rate increases linearly with DNA length while the off-rate is independent of length for DNA longer than 60 bp. Next, we used an optical trap equipped with a scanning confocal microscope to show that SWR1 can diffuse one-dimensionally along stretched DNA, with a diffusion coefficient that permits scanning of a typical NDR in 93 milliseconds. Interestingly, we see that ATP binding alone increases the one-dimensional diffusion coefficient of SWR1 along DNA. We found that a major DNA binding subunit of the SWR1 complex, Swc2, also diffuses on DNA suggesting that it contributes to SWR1’s diffusivity. The diffusion coefficient for both SWR1 and Swc2 increases with ionic strength suggesting that SWR1 utilizes some microscopic dissociation and reassociation events, known as hopping, to diffuse on DNA. However, it is likely that SWR1 only makes infrequent hops, with most of the diffusion on DNA being mediated by helically coupled diffusion, known as sliding, since SWR1 diffusion is blocked by proteins that are bound to DNA, such as dCas9, and the diffusion of the complex is slower than would be expected for majority hopping diffusion. Lastly, we observed SWR1 diffusion on DNA containing sparsely deposited nucleosomes and found that SWR1 diffusion is confined between nucleosomes. Our data indicates that a multi-subunit chromatin remodeler can diffuse along DNA and suggests that SWR1 finds its target, the +1 nucleosome, through facilitated diffusion. Facilitated diffusion may be a common search mechanism for all chromatin remodelers that act upon nucleosomes positioned next to free DNA, such as those adjacent to the NDR.

## Results

### SWR1 binding kinetics depend on DNA length

To study both the DNA binding kinetics and diffusive behavior of SWR1, we generated a site-specifically labeled complex referred to as Cy3-SWR1 (**Figure 1B**). We purified SWR1 from *S. cerevisiae* in the absence of the Swc7 subunit (SWR1ΔSwc7). Recombinant Swc7 was expressed and purified from *E. coli*, a single cysteine in Swc7 was labeled with Cy3, and the labeled Swc7 was then added to the SWR1ΔSwc7 preparation between two steps of a specialized tandem affinity purification protocol(Sun et al., 2020). Subsequent purification on a glycerol gradient revealed that the Cy3-labeled Swc7 co-migrated with the rest of the SWR1 subunits, demonstrating incorporation of Swc7 back into the SWR1 complex (**Figure 1–figure supplement 1A**). The histone exchange activity of the labeled Cy3-SWR1 was identical to that of wild type SWR1 as revealed by an electrophoretic mobility shift assay (EMSA) (**Figure 1–figure supplement 1B**).

While it is well established that the affinity of SWR1 for DNA is dependent on DNA length (Ranjan *et al*., 2013), the kinetics of binding are unknown. We used single-molecule colocalization measurements to observe Cy3-SWR1 binding and unbinding on Cy5-labeled DNA of different lengths in real time (**Figure 1C-E**). These measurements showed that both the on-rate (*k*_bind_) and the lifetime of the SWR1-DNA complex (*t*_off_) are dependent on DNA length. The on-rate for SWR1 binding to 20 bp DNA, the approximate size of linker DNA between intragenic nucleosomes in yeast, was 1×10^6^ M^-1^ s^-1^. Increasing the DNA length to 150 bp, the approximate size of the NDR in yeast, increases the binding rate 36-fold to 3.6×10^7^ M^-1^ s^-1^. *k*_bind_ increased linearly with DNA length between these two values (**Figure 1F**). Interestingly, we found that DNA could accommodate multiple bound SWR1 molecules, with the likelihood of multiple binding events increasing with DNA length (see **Figure 1E** for example trace). Cy3-Swc7 alone exhibited no affinity for 150 bp DNA (data not shown), suggesting that the observed Cy3-signal increase is caused by the full Cy3-SWR1 complex binding to DNA.

The lifetime of SWR1 bound to DNA (*t*_off_) was also sensitive to DNA length, exhibiting two sharp increases as DNA size increased from 20 to 40 bp, and 60 to 80 bp. Whereas *t*_off_ for 20 bp DNA was 1.5 +/- 0.3 s, *t*_off_ for SWR1 binding to 40 and 60 bp DNA increased to 9 +/- 1.4 s and 12 +/- 5.8 s, respectively, which is the same within error (**Figure 1G**). Once the DNA was 80 bp or longer, however, the lifetime increased dramatically to at least 30 s, which is the photobleaching limit of the measurement (**Figure 1–figure supplement 1C**). Measurements at low laser power showed that SWR1 remained bound to 150 bp DNA for at least 5 minutes. *t*_off_ was unchanged in the presence of ATP but was sensitive to ionic strength, decreasing with added salt (**Figure 1D-E**). Curiously, *t*_off_ also decreased in the presence of competitor DNA (**Figure 1–figure supplement 1D-E**). The kinetic measurements show that the affinity of SWR1 for DNA greater than 60 bp is primarily limited by the on-rate, suggesting the increased occupancy of SWR1 at longer NDRs observed in yeast (Ranjan *et al*., 2013) is a result of the increased probability of SWR1 finding the NDR, as opposed to an increase in the residence time of SWR1.

### SWR1 scans DNA

To determine if SWR1 can move along DNA, we tracked single Cy3-SWR1 complexes bound to stretched lambda DNA using an optical trap equipped with a confocal scanning microscope (LUMICKS, C-Trap) (Heller et al., 2014a; Heller et al., 2014b). The experiment was carried out using a commercial flow-cell in order to efficiently catch beads, trap DNA, and image bound proteins over time (**Figure 2A**) as has been performed previously (Balaguer et al., 2021; Brouwer et al., 2016; Gutierrez-Escribano et al., 2019; Newton et al., 2019; Rill et al., 2020; Wasserman et al., 2019). Briefly, lambda DNA end-labeled with biotin is tethered between two optically trapped streptavidin-coated polystyrene beads, pulled to 5 piconewton (pN) tension to straighten the DNA (Baumann et al., 2000) and the distance between the two optical traps is clamped (**Figure 2A-B**). After confirming the presence of a single DNA tether, the DNA is brought into an adjacent channel of the flow-cell containing 250 picomolar Cy3-SWR1. Confocal point scanning across the length of the DNA was used to image single Cy3-SWR1 bound to lambda DNA over time to generate kymographs (**Figure 2B-C**). The observed fluorescent spots represent the Cy3-SWR1 complex as Cy3-Swc7 alone was unable to bind DNA (**Figure 2–figure supplement 1**).

**Fig. 2.**
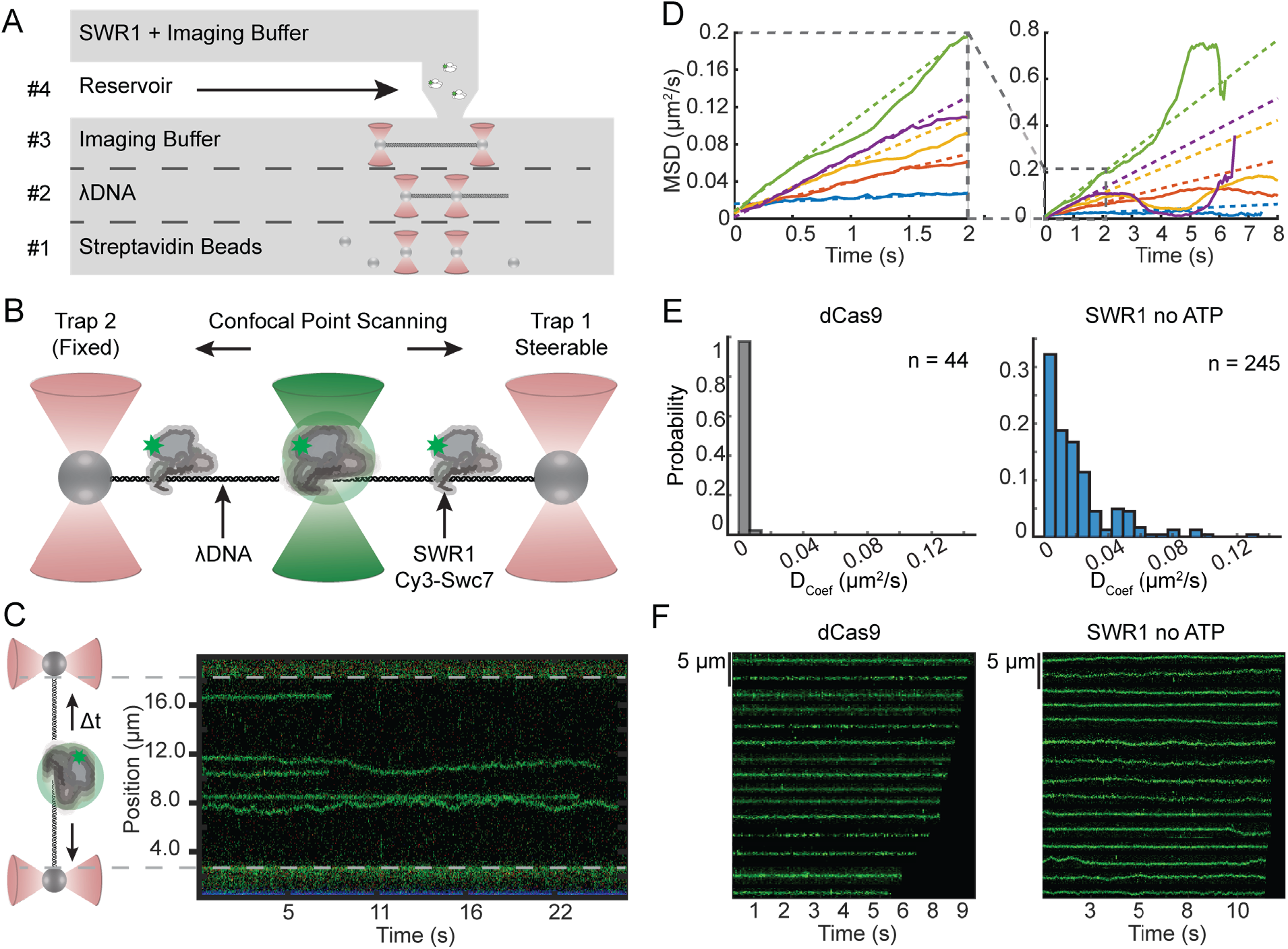
SWR1 diffuses on extended dsDNA. (A) Schematic representation of a C-Trap microfluidics imaging chamber with experimental workflow depicted therein: #1 catch beads, #2 catch DNA, #3 verify single tether, #4 image SWR1 bound to DNA. (B) Schematic representation of confocal point scanning across the length of lambda DNA tethered between two optically trapped beads. This method is used to monitor the position of fluorescently labeled SWR1 bound to DNA. (C) Example kymograph with a side-by-side schematic aiding in the interpretation of the kymograph orientation. (D) Mean squared displacement (MSD) versus time for a random subset of SWR1 traces in which no ATP is added. An enlargement of the initial linear portion is shown to the left where colored dashed lines are linear fits to this portion. (E) Histogram of diffusion coefficients for dCas9 (left) and SWR1 in which no ATP is added (right) (F) Segmented traces of dCas9 (left) and SWR1 in which no ATP is added (right). **Figure 2 – Source data 1** Data underlying panel D and E **Figure 2 – Source data 1** Uncropped kymograph Tiff image from panel C

Cy3-SWR1 bound to lambda DNA is mobile, demonstrating that Cy3-SWR1 can move on DNA once bound and the movement did not appear to be unidirectional. Therefore, we plotted mean square displacement (MSD) vs time and found that the initial portion of the curve is linear, suggesting that diffusion is Brownian (**Figure 2D**). The diffusion coefficient observed (D_1,obs_) for Cy3-SWR1 was 0.013±0.002 μm^2^/sec in buffer alone (**Figure 2E-F**). Since diffusion coefficient distributions are non-normal, D_1,obs_ is defined as the median diffusion coefficient of all molecules in a condition; individual diffusion coefficients were determined from the slope of the initially linear portion of their respective MSD plot (see **Materials and Methods** for more details). This diffusion coefficient is comparable to other proteins with characterized 1D diffusion(Gorman et al., 2007; Park et al., 2021). In contrast D_1,obs_ for specifically bound Cy5-dCas9, an immobile reference with D_1,obs_ of 0.0003±0.0004 μm^2^/sec, is forty times smaller than Cy3-SWR1. These measurements clearly show that SWR1 undergoes Brownian diffusion on nucleosome-free DNA.

### ATP bound SWR1 is more diffusive than the unbound complex

To determine if SWR1 can actively translocate on DNA, we observed the motion of Cy3-SWR1 in the presence of 1 mM ATP (**Figure 3**). The MSDs of Cy3-SWR1 in the presence of ATP remained linear, showing that SWR1 does not translocate directionally on DNA (**Figure 3A**). The increased slope of the MSDs in the ATP condition, however, does indicate that ATP increases the diffusion. An overlay of all trajectories from both conditions further demonstrates that SWR1 diffuses a greater distance from the starting position in the presence of ATP and that its motion is not directional (**Figure 3B**). To address whether this increased diffusion was due to ATP hydrolysis, we also measured SWR1 diffusion in the presence of 1 mM ATP*γ*S, a nonhydrolyzable analog of ATP, as well as with ADP. The distribution in diffusion coefficients in the presence of ATP and ATP*γ*S are both shifted to higher values compared to in the absence of ATP or in the presence of ADP (**Figure 3C**). This shift was shown to be statistically significant using the non-parametric Mann-Whitney U-test (**Figure 3D**). SWR1 diffusion in the presence of 1mM ATP (D_1,obs =_ 0.024 μm^2^/sec ± 0.001) was not significantly different than diffusion in the presence of 1 mM ATP*γ*S (D_1,obs =_ 0.026 μm^2^/sec ± 0.002). Similarly, SWR1 diffusion in the absence of ATP (D_1,obs =_ 0.013 μm^2^/sec ± 0.002) was not different than SWR1 diffusion in the presence of 1mM ADP (D_1,obs =_ 0.011 μm^2^/sec ± 0.002). Additionally, we found that ATP decreased the fraction of slow or immobile Cy3-SWR1 molecules, defined as those molecules that show D_1_ values that are indistinguishable from dCas9 values (**Figure 3E**). While 9% of Cy3-SWR1 were slow or immobile in the presence of ATP, 32% were slow or immobile in buffer alone. These results show that while SWR1 does not actively translocate on DNA, binding of ATP increases the mobility of SWR1 on DNA.

**Fig. 3.**
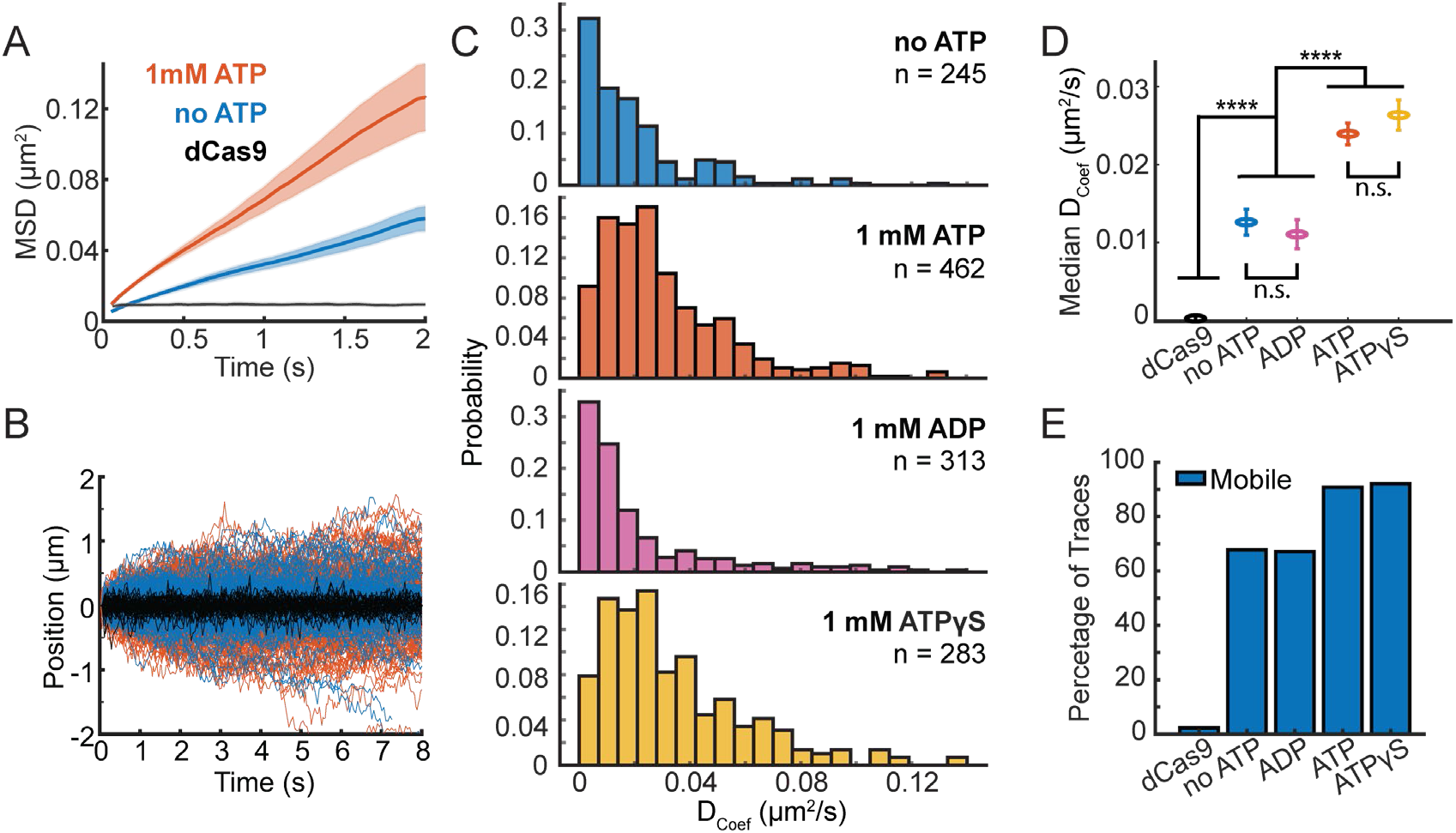
ATP binding modulates SWR1 diffusion. (A) Mean MSD vs time plotted for 1mM ATP (orange, n= 124), no ATP (blue, n = 134), and dCas9 (black, n = 25) with shaded error bars SEM. (B) SWR1 trajectories aligned at their starts for 1mM ATP (orange lines), no ATP (blue lines), and dCas9 as reference for immobility (black lines). All trajectories represented. (C) Histograms of diffusion coefficients extracted from individual trajectories for SWR1 diffusion in the presence of no ATP, 1mM ATP, 1mM ADP, 1mM ATP*γ*S (from top to bottom). The number of molecules measured (n) for each condition is printed in each panel. (D) Median diffusion coefficients for SWR1 in varying nucleotide conditions. dCas9 is shown as a reference. Error bars are the uncertainty of the median. (E) Percentage of mobile traces in each condition, where immobility is defined as traces with similar diffusion coefficients to dCas9 (defined as diffusion coefficients smaller than 0.007 µm^2^/sec). **Figure 3 – Source data 1** Numerical data underlying panels A, C, D, and E

### SWR1 and the DNA binding domain of the Swc2 subunit slide on DNA

SWR1 binding to DNA is mediated in part by the Swc2 subunit, which harbors a positively charged and unstructured DNA binding domain (Ranjan *et al*., 2013). To determine if Swc2 contributes to the diffusive behavior of SWR1 on DNA we compared diffusion coefficients of the SWR1 complex to diffusion coefficients of the DNA binding domain (DBD) of Swc2 (residues 136-345, **Figure 4–figure supplement 1**). We found that Swc2 also diffuses on DNA, however the median diffusion coefficient, D_1,obs =_ 1.04 μm^2^/sec ± 0.09, was approximately 40-fold larger than that of SWR1 in the presence of 1mM ATP (**Figure 4, Materials and Methods**). This large difference in measured diffusion coefficients could be due to the difference in size between the small Swc2 DBD and full SWR1 complex or to other DNA binding components of SWR1 interacting with DNA and increasing friction. Based on theoretical models of rotation coupled versus uncoupled diffusion, the scaling relationship between size and diffusion coefficient is consistent with SWR1 and Swc2 DBD utilizing rotationally-coupled sliding(Blainey et al., 2009) (**Figure 4–figure supplement 2**).

**Fig. 4.**
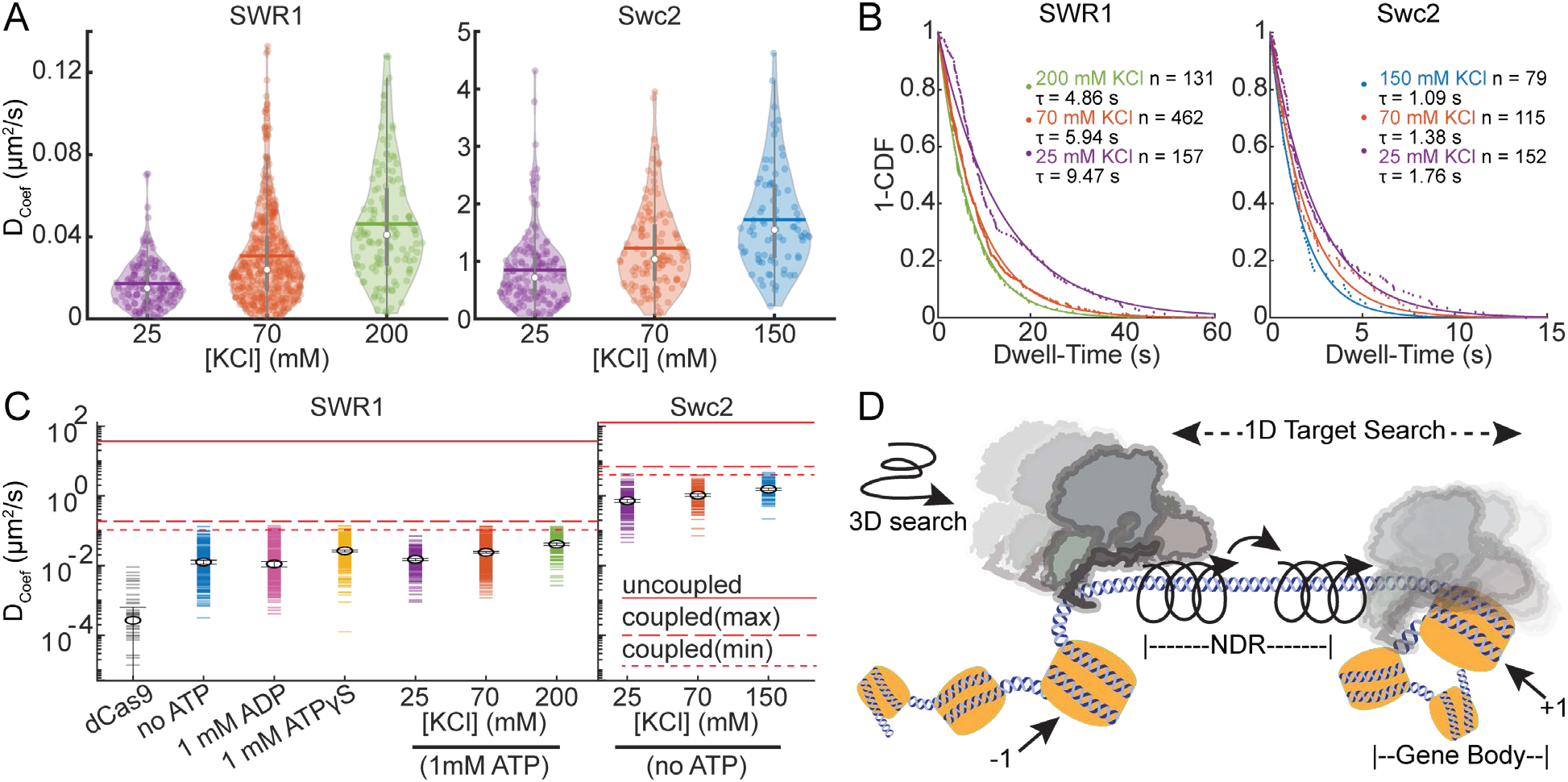
SWR1 and Swc2 DBD utilize sliding to scan DNA. (A) Violin plots of diffusion coefficients for SWR1 and Swc2 DNA binding domain (DBD) in increasing potassium chloride concentrations. Medians are shown as white circles and the mean is indicated with a thick horizontal line. (B) 1-CDF plots of SWR1 and Swc2 were fit to exponential decay functions to determine half-lives of binding in varying concentrations of potassium chloride. The number of molecules as well as half-lives determined are printed therein. Dots represent data points, while solid lines represent fits. Half-lives are calculated using the length of all the trajectories in each condition. (C) Upper limits for diffusion of SWR1 and Swc2 predicted using either a helically uncoupled model for hopping diffusion (uppermost solid red line) or a helically coupled model for sliding diffusion (lower dashed red lines). Two dashed lines are shown for helically coupled upper limits because the distance between the helical axis of DNA and the center of mass of either SWR1 or Swc2 is unknown. Markers represent median values. D_coef_ values for each condition are shown as horizontal dashes, the number of molecules represented in each condition is as aforementioned. (D) A schematic representation of a model for how SWR1 likely performs 1D diffusion on DNA. **Figure 4 – Source data 1** Data underlying panels A, B, and C

Next, we found that both SWR1 and Swc2 DBD show increased diffusion with increasing concentrations of potassium chloride (**Figure 4A)**, and each showed decreasing binding lifetimes with increasing salt (**Figure 4B**). Both increased diffusion and decreased binding lifetimes are features of 1D hopping, as the more time a protein spends in microscopic dissociation and reassociation the faster it can move on DNA, but also falls off DNA more frequently(Bonnet et al., 2008; Mirny et al., 2009). This data is consistent with the single molecule TIRF data presented earlier (**Figure 1–figure supplement 1E**), which also reveals decreased binding lifetimes to DNAs when ionic strength is increased. The TIRF assay also shows that competitor DNA can decrease binding lifetime as would be expected for a protein that hops on DNA and may be prone to alternative binding onto competitor DNA (Brown et al., 2016; Gorman *et al*., 2007).

The theoretical upper limit of diffusion for a particle that uses linear translocation (1D hopping) is higher than the theoretical upper limit of diffusion with helically coupled sliding because in the latter there are additional rotational components of friction incurred when circumnavigating the DNA axis (Blainey *et al*., 2009). Based on the molecular weight of SWR1 and Swc2, the theoretical upper limits of 1D diffusion using rotation coupled versus uncoupled 1D diffusion can be calculated (**Materials and Methods**). In all conditions measured, the median diffusion of SWR1 is below the upper limit with rotation (**Figure 4C**), consistent with much of the observed diffusion coming from SWR1 engaging in rotationally coupled diffusion. Nonetheless, some individual traces have diffusion coefficients that surpass this theoretical maximum, indicating that there may be alternative modes for engaging with DNA (e.g., infrequent hopping), which allows it to surpass the upper limit with rotation (Ahmadi et al., 2018; Gorman et al., 2010). A similar phenomenon was observed for Swc2 DBD, which also exhibited median diffusion coefficients below the theoretical maximum with rotation, with some traces having diffusion coefficients above this limit (**Figure 4C**). These trends are consistent with a model in which SWR1 utilizes a majority 1D helically coupled sliding with occasional hopping to diffuse on DNA (**Figure 4D**).

### SWR1 cannot bypass bound dCas9

While the nucleosome depleted region is a region of open chromatin where accessibility to DNA is higher compared to DNA in gene bodies, SWR1 must compete with transcription factors and other DNA binding proteins for search on this DNA (Kim *et al*., 2021; Kubik et al., 2019; Nguyen et al., 2021; Rhee and Pugh, 2012). Proteins that diffuse on DNA by 1D hopping are sometimes capable of bypassing protein barriers and nucleosomes (Gorman *et al*., 2010; Hedglin and O’Brien, 2010). To investigate whether ATP bound SWR1 can bypass protein barriers, we turned to dCas9, an endonuclease inactive mutant of Cas9, to serve as a programmable barrier to diffusion. We used a dual color single particle tracking scheme to simultaneously observe Cy3-labeled SWR1 diffusion and the positions of Cy5-labeled dCas9 (**Figure 5**). crRNAs were used to direct dCas9 binding to 5 positions on the lambda DNA using previously validated targeting sequences (**Figure 5A, Table 2, Materials and Methods**) (Sternberg *et al*., 2014). We assume that dCas9 binding far outlasts the photobleaching lifetime of Cy5 (Singh et al., 2016), therefore we use the average position of the particle to extend the trace after photobleaching of Cy5 for colocalization analysis. Out of 106 traces with colocalization events, 67% showed SWR1 moving away from dCas9 toward where it came from as if it was reflected from a boundary (**Figure 5B, C**). Another 30% of traces showed SWR1 immobile and colocalized with dCas9 for the duration of the trace, which we describe as stuck (**Figure 5C, D**). Only 3% of all colocalization events exhibited a cross-over event (**Figure 5C, E, Figure 5–figure supplement 1)**. The ability of dCas9 to block SWR1 diffusion in most encounters further supports a model in which SWR1 mainly engages in helically coupled sliding (**Figure 4D**). Infrequent hopping events that colocalize to a dCas9 encounter may contribute to the presence of the rare bypass event (**Figure 5E, Figure 5–figure supplement 1)**.

**Fig. 5.**
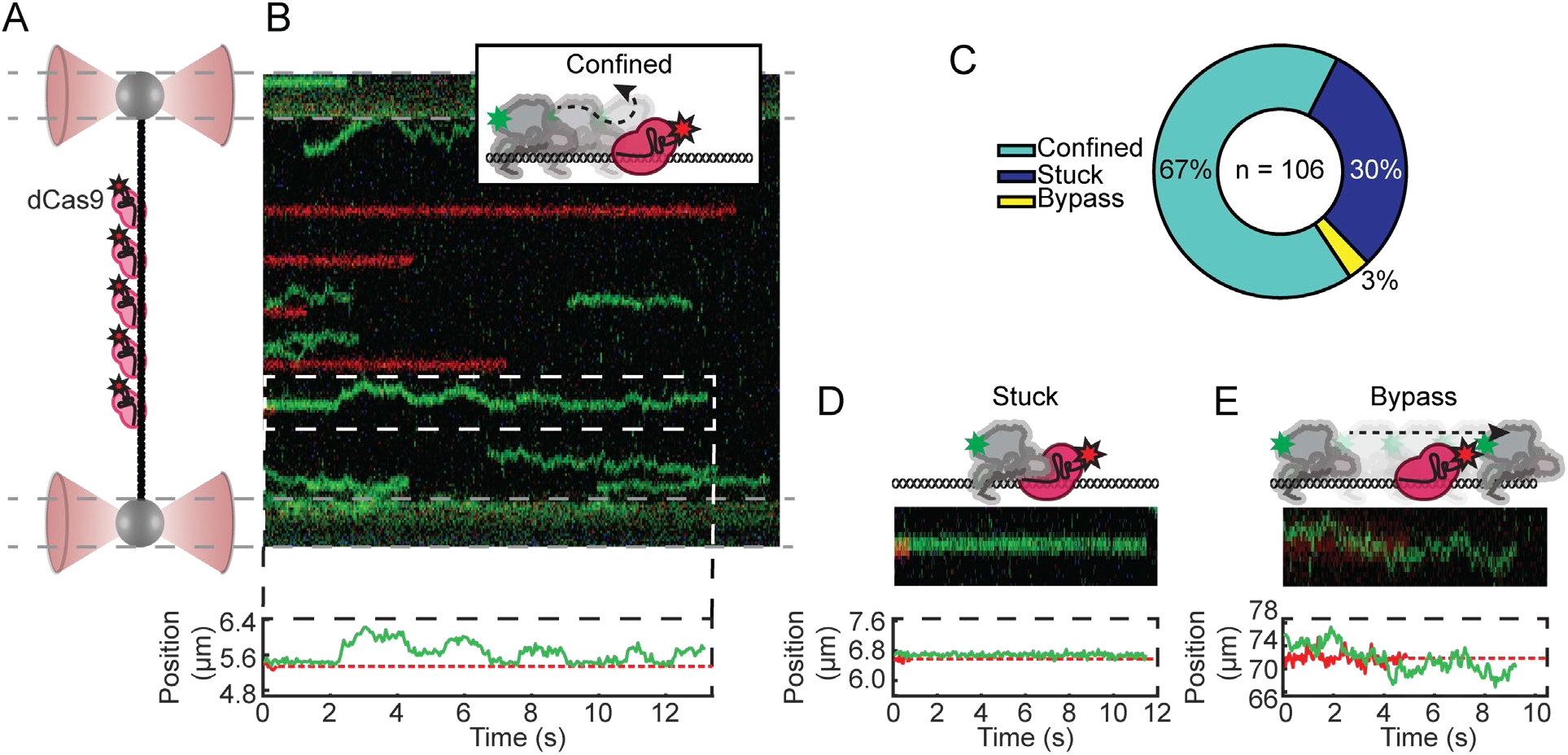
SWR1 protein roadblock bypass assay. (A) Schematic of the experimental set-up: 5 Cy5-labeled gRNA position dCas9 at 5 evenly spaced sites along lambda DNA. (B) Example kymograph with 5 bound dCas9 in red, and an example of a confined diffusion encounter. Schematic, and single particle tracking trajectory printed above and below. (C) Pie-chart of the three types of colocalization events with the total number of observations printed therein. (D) Example of SWR1 stuck to the dCas9 within limits of detection; schematic, cropped kymograph, and single particle tracking trajectory shown. (E) Example of a SWR1-dCas9 bypass event; schematic, cropped kymograph, and single particle tracking trajectory shown. (B, D, E) In the example single particle tracking trajectory, dCas9 is represented as a dashed red line after Cy5 has photobleached, however due to long binding lifetimes of dCas9 we continue to use its position for colocalization analysis. **Figure 5 – Source data 1** All colocalization events with classifications indicated

### Nucleosomes are barriers to SWR1 diffusion

Diffusion over nucleosomes may also be an important aspect of target search; it is not known whether SWR1 diffusing on an NDR would be confined to this stretch of DNA by flanking +1 and -1 nucleosomes or whether its diffusion could continue into the gene body (**Figure 6A**). To investigate this, we monitored SWR1 diffusion on sparse nucleosome arrays reconstituted on lambda DNA. Nucleosomes were formed at random sites along lambda DNA using salt gradient dialysis, as has been done previously (Gruszka et al., 2020; Visnapuu and Greene, 2009) (**Figure 6–figure supplement 1, Materials and Methods**). On average, 40 ± 5 nucleosomes were incorporated onto the lambda nucleosome arrays as shown by nucleosome unwrapping force-distance curves (**Figure 6B-E**); nucleosomes showed detectable unwrapping at forces 15 pN or greater (**Figure 6B**) (Brower-Toland et al., 2002; Fierz and Poirier, 2019), with a characteristic lengthening of the array by ∼25 nm with each unwrapping event (**Figure 6C**)(Spakman et al., 2020). To determine a compaction ratio which could be used to estimate the number of nucleosomes per array in the case where the array breaks before it has been fully unwrapped, unwrapping events were counted and related to the total length of the array at 5pN (**Figure 6–figure supplement 1A, Materials and Methods**).

**Fig. 6.**
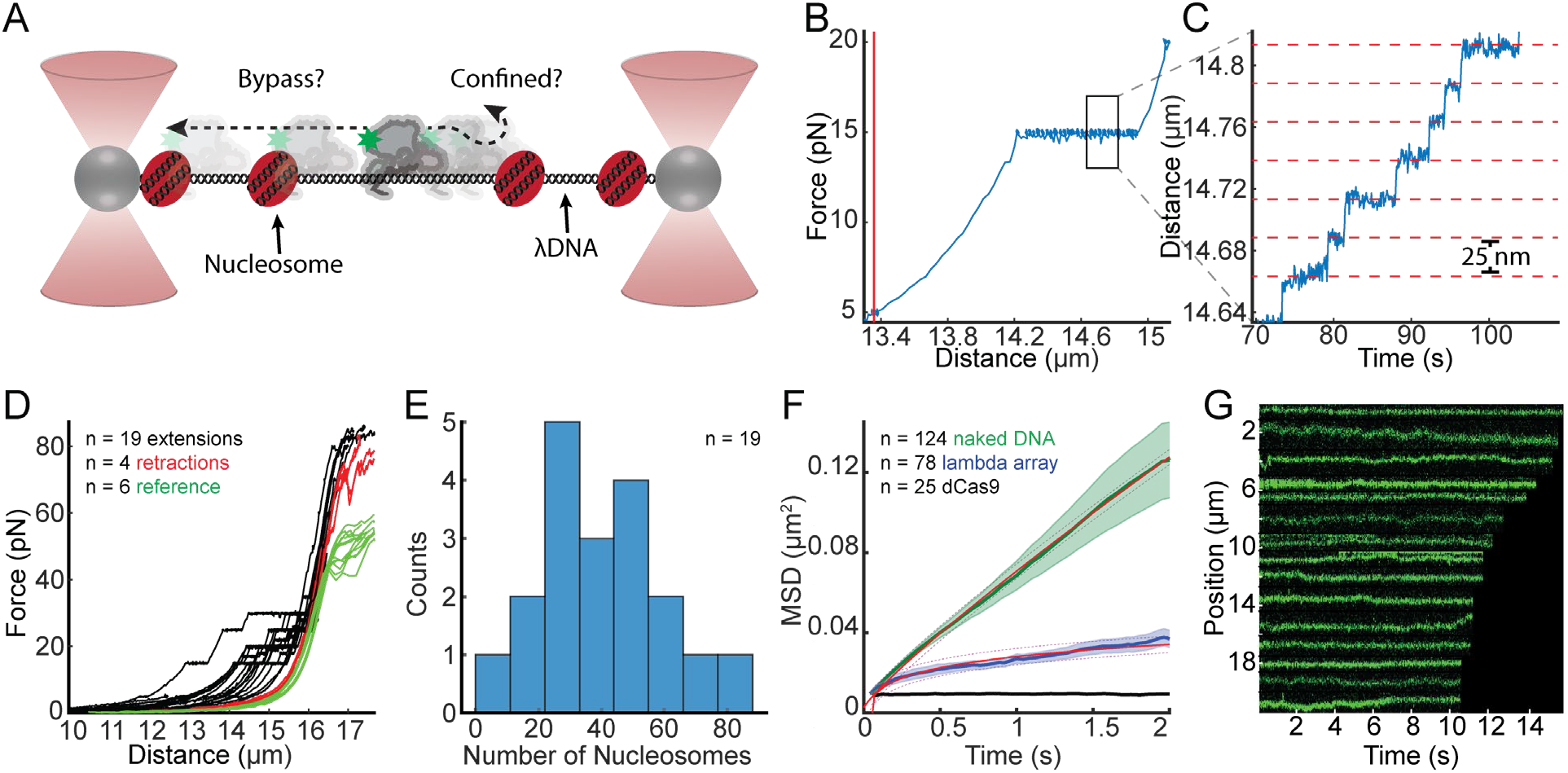
SWR1 does not diffuse over nucleosomes. (A) Schematic of the experimental set-up, with experimental question depicted therein. (B) Example force-distance curve showing that at 15pN nucleosomes begin to unwrap. Vertical red line shows the length of the nucleosome array 5pN. (C) Example unwrapping events that result in characteristic lengthening of 25 nm at this force regime. (D) Lambda nucleosome arrays extension (unwrapping) and retraction curves, with a reference naked DNA force-distance curves. Black curves are unwrapping curves where the force is clamped at either 20, 25 or 30 pN to visualize individual unwrapping events; red curves are the collapse of the DNA after unwrapping nucleosomes; green curves are reference force extension plots of lambda DNA without nucleosomes. (E) Histogram of the number of nucleosomes per array determined from the length of the array at 5pN and the compaction ratio. (F) Mean MSDs are fit over the first 2 seconds to MSD = Dt^α^, the red lines represent the fits with 95% confidence interval shown as dashed lines. SWR1 diffusing on naked DNA [green curve], on lambda nucleosome arrays [blue] and for comparison dCas9 [black]. (G) Representative SWR1 particles diffusing on the nucleosome arrays are cropped and arranged by the length of the trace. **Figure 6 – Source data 1** Data underlying panels B, C, and E

Overall, the behavior of SWR1 on lambda nucleosome arrays was notably different than on naked lambda DNA (**Figure 6F-G**). The mean MSD for SWR1 on naked DNA increases linearly at short time scales (< 2 s), whereas the mean MSD for SWR1 on the lambda nucleosome array plateaus over this same time scale, indicative of confined 1D diffusion (**Figure 6F**). The degree to which diffusion is confined can be described by α<1 where MSD = Dt^α^. Whereas SWR1 on naked DNA has an α = 0.88 over a 2 second time scale, SWR1 on the lambda array has an α = 0.24 reflecting considerable confinement. By fitting the MSD curve to an exponential function, the mean MSD appears to approach a limit of 0.054 μm^2^ (**Figure 6–figure supplement 1B**). Assuming an even distribution of an average of 40 nucleosomes per array (**Figure 6B**), the mean distance between nucleosomes is equal to 0.35 μm; whereas the length of DNA to which SWR1 diffusion is confined is approximately 0.23 μm, determined from the square root of the MSD limit described above. Representative traces show signs of confinement, as more immobile segments dominate the trace and the range of SWR1 exploration becomes confined (**Figure 6G**). The data, therefore, suggests that SWR1 diffusion is confined to the space between nucleosomes.

## Discussion

### Reducing the dimensionality of nucleosome target search

Our single molecule tracking data shows that SWR1 slides on DNA, which is a novel finding for a chromatin remodeler. Moreover, SWR1 scans DNA with a diffusion coefficient comparable to other well-characterized proteins that utilize facilitated diffusion to bind specific DNA sequences or lesions (Ahmadi *et al*., 2018; Blainey et al., 2006; Gorman *et al*., 2010; Kamagata et al., 2020; Porecha and Stivers, 2008; Tafvizi et al., 2011; Tafvizi et al., 2008; Vestergaard et al., 2018). Without 1D sliding, the search process of SWR1 for its target nucleosome would be dependent solely on 3D collisions with nucleosomes. In the yeast genome, there are approximately 61,568 annotated nucleosomes (Jiang and Pugh, 2009; Kubik et al., 2015), of which 4,576 are identified as potential +1 nucleosomes enriched in H2A.Z (Tramantano et al., 2016). Since only 7% of nucleosomes are targets of SWR1 histone exchange, we believe that +1 nucleosomes use their adjacent NDRs as antennas, promoting SWR1 binding and 1D search to encounter flanking nucleosomes (Mirny *et al*., 2009). This increased efficiency in target localization through dimensional reduction of the search process may be one that could extend to other chromatin remodelers that act on nucleosomes adjacent to the NDR, such as RSC, SWI/SNF, CHD1, ISW1, ISW2, and INO80 (Kim *et al*., 2021).

### ATP binding facilitates SWR1 target search and diffusion on DNA

We observed that SWR1 diffusion is increased in the presence of ATP, and that substitution with ATP*γ*S also results in similar increased diffusion suggesting that this enhancement is mediated by nucleotide binding rather than hydrolysis. SWR1 requires ATP to perform the histone exchange reaction, and basal levels of ATP hydrolysis when any one of the required substrates for the histone exchange reaction is missing is low (Luk et al., 2010). This includes the scenario where SWR1 is bound to DNA in the absence of the nucleosome and H2A.Z/H2B dimer. Therefore, we do not expect SWR1 diffusion in the presence of 1mM ATP to be modulated by ATP hydrolysis, which is consistent with our findings. Binding of nucleotide cofactor has been shown to produce conformational changes in ATPases that can affect their diffusion on DNA (Gorman *et al*., 2007). The core ATPase domain of SWR1, Swr1, like other chromatin remodelers, belongs to the superfamily 2 (SF2) of translocases which have two lobes that switch between an open and closed conformation with ATP binding and hydrolysis (Beyer et al., 2013; Nodelman et al., 2020). It is therefore possible that the ATP bound closed conformation of the core ATPase results in a DNA binding interface, distributed across accessory domains, that is more conducive to diffusion on DNA, contributing to the enhanced diffusion of SWR1 in the presence of ATP or ATP*γ*S. In the present study we further investigated SWR1’s main DNA binding subunit, Swc2, which forms an extended interface with the core ATPase (Willhoft et al., 2018). In addition to the changes in the contacts that the translocase domain makes with DNA in the closed versus open form, it is possible that ATP modulates how Swc2 engages with the DNA through conformational changes propagated from Swr1. Swc2 appears to be an important accessory subunit for 1D diffusion, as we were able to show that in isolation, the DNA binding domain of Swc2 slides on DNA with properties similar to that of the whole complex although with a much-increased diffusion coefficient.

Conformations that result in slower sliding presumably become trapped in free energy minima along the DNA where the DNA sequence or the presence of DNA lesions results in a more stably bound DNA-protein interaction (Gorman *et al*., 2007). While it remains unknown whether SWR1 interacts with different sequences of DNA differently in the context of sliding, we believe this may be a possibility since we observe a distribution in diffusion coefficients within any single condition which would not be expected if the energetic costs of binding substrate were equal everywhere. The NDR is rich in AT-content; therefore one might imagine that SWR1 may have evolved to be better at scanning DNA with high AT-content (Chereji et al., 2018). Lambda DNA, the DNA substrate used in this study, has asymmetric AT-content, which has been shown to affect nucleosome positioning during random deposition (Visnapuu and Greene, 2009). Future studies of chromatin remodeler 1D diffusion are needed to address this possibility.

### SWR1 and Swc2 predominantly slide with diffusion confined between roadblocks

The way a protein engages with DNA during 1D search can have impacts on both scanning speed and target localization. For instance, a protein that maintains continuous contact with the DNA in part through charge-charge interactions with the phosphate backbone will predominantly utilize helically coupled sliding. By contrast, a protein that dissociates just far enough from the DNA for cation condensation on the phosphate backbone to occur before quickly reassociating will utilize linear hopping to perform short 3D searches before reassociating at a nearby site on the DNA (Mirny *et al*., 2009). Proteins that hop on DNA therefore have increased diffusion with increased monovalent cation concentration, as a higher screening potential results in more frequent hops. SWR1 and the DNA binding domain of the Swc2 subunit both become more diffusive as the concentration of potassium chloride is increased (**Figure 4A**), which indicates that both utilize some degree of hopping when diffusing on DNA.

Nonetheless, the observed diffusion for both SWR1 and Swc2, on average, falls within a range expected for a protein that predominantly uses a sliding mechanism to diffuse on DNA. In order for a protein to slide or hop on DNA, the energy barrier (ΔG^‡^) to break the static interaction and dynamically engage with the DNA following the parameters of either the sliding or hopping model must be less than ≈ 2 k_B_T (Ahmadi *et al*., 2018; Gorman *et al*., 2007; Slutsky and Mirny, 2004). Based on the molecular weight of SWR1 and Swc2, the upper limit of 1D diffusion was estimated for both the sliding and hopping model (**Figure 4C, Materials and Methods)**. The upper limit of diffusion coefficients for rotation-coupled sliding-only diffusion is lower than hopping-only diffusion due to the rotational component increasing friction in the sliding model. We found that most particles for either SWR1 or Swc2 fall below the estimated upper limit for sliding diffusion. This observation indicates that, averaged over the length of the trace, the energetic barrier to exclusively hop along DNA is too large, whereas the energy barrier for sliding diffusion is permissive (<2 k_B_T). Therefore, while both SWR1 and Swc2 DNA binding domain can engage in hopping, both on average utilize sliding diffusion as exhibited by their slow diffusion.

Sliding as a predominant component of the SWR1 interaction with DNA is further evidenced by the observation that SWR1 can neither bypass a dCas9 protein roadblock nor nucleosomes with high efficiency. Other studies have found that proteins that utilize sliding as the predominant form of 1D diffusion cannot bypass proteins or nucleosomes (Brown *et al*., 2016; Gorman *et al*., 2010; Hedglin and O’Brien, 2010), whereas a protein that predominantly hops may be able to bypass these obstacles. The utilization of hopping diffusion has been described as a trade-off between scanning speed and accuracy, with proven implications in target sequence bypass by the transcription factor LacI (Marklund et al., 2020). Whether the same may be true for chromatin remodelers in search of specific nucleosomes is yet to be reported.

### Concluding remarks

Single particle tracking in vivo has shown that approximately 47% of SWR1 molecules are bound to chromatin and the remainder is performing 3D diffusion (Ranjan et al., 2020). Once bound (e.g. near the center of an average NDR of ∼150 bp) our findings suggest that SWR1 would require 46 milliseconds (see **Materials and Methods**) to scan and encounter a flanking nucleosome by 1D diffusion at 0.024 μm^2^/sec. A recent report shows that when complexed with a canonical nucleosome and the H2A.Z-H2B dimer, SWR1 can rapidly perform the ATP hydrolysis-dependent histone exchange reaction, which occurs on average in 2.4 seconds as measured by an in vitro single molecule FRET assay (Poyton et al., 2021). Thus, SWR1-catalyzed histone H2A.Z exchange on chromatin may be an intrinsically rapid event that occurs on a timescale of seconds. While 1D diffusion should in principle allow SWR1 to encounter either the +1 or -1 nucleosome at the ends of the NDR, directionality may be conferred by the preferentially acetylated +1 nucleosome, where interaction with SWR1’s two bromodomains on the Bdf1 subunit should increase binding lifetime during encounter events (Ranjan et al., 2013). Future studies of 1D diffusion with the use of nucleosome arrays that mimic the natural nucleosome arrangement and histone modifications of NDRs and gene bodies should provide important physical and temporal insights on how SWR1 undergoes target search to capture its nucleosome substrates at gene promoters and enhancers. Extension of this approach to other ATP-dependent chromatin remodelers and histone modification enzymes will facilitate understanding of the cooperating and competing processes on chromatin resulting in permissive or nonpermissive architectures for eukaryotic transcription.

## Materials and Methods

### Protein purification, fluorescence labeling, and functional validation (SWR1 & Swc2)

The SWR1 complex labeled only on Swc7 was constructed as has been previously documented (Poyton *et al*., 2021). We demonstrated that the fluorescently labeled SWR1 complex maintains full histone exchange activity (**Figure 1–figure supplement 1B**). For this assay, 1 nM SWR1, 5 nM nucleosome, and 15 nM ZB-3X flag were combined in standard SWR1 reaction buffer [25 mM HEPES pH 7.6, 0.37 mM EDTA, 5% glycerol, 0.017% NP40, 70 mM KCl, 3.6 mM MgCl_2_, 0.1 mg/mL BSA, 1 mM BME] supplemented with 1 mM ATP, and the reaction was allowed to proceed for 1 hour before being quenched with (100 ng) lambda DNA. The product was run on a 6% native mini-PAGE run in 0.5X TB as has been previously reported (Ranjan *et al*., 2013).

The DNA binding domain (DBD) of Swc2 (residues 136-345) was cloned into a 6x his-tag expression vector with a single cysteine placed directly before the N-terminus of the protein for labeling purposes (**Table 1**). The Swc2 DBD was purified after expression under denaturing conditions using Ni-NTA affinity purification. After purification, the Swc2 DBD was specifically labeled in a 30-fold excess of Cy3-maleimide. After fluorophore labeling the Swc2 DBD was Ni-NTA purified a second time to remove any excess free dye. The product was then dialyzed overnight at 4°C into refolding buffer [20 mM Tris pH 8.0, 0.5 M NaCl, 10% Glycerol, 2 mM β-Mercaptoethanol, 0.02% NP40 and 1 mM PMSF] as has been previously documented (Ranjan *et al*., 2013). Pure protein was stored as aliquots at - 80°C until time of use. SDS-page reveals a pure Cy3-labeled product (**Figure 4–figure supplement 1**).

**Table 1.**
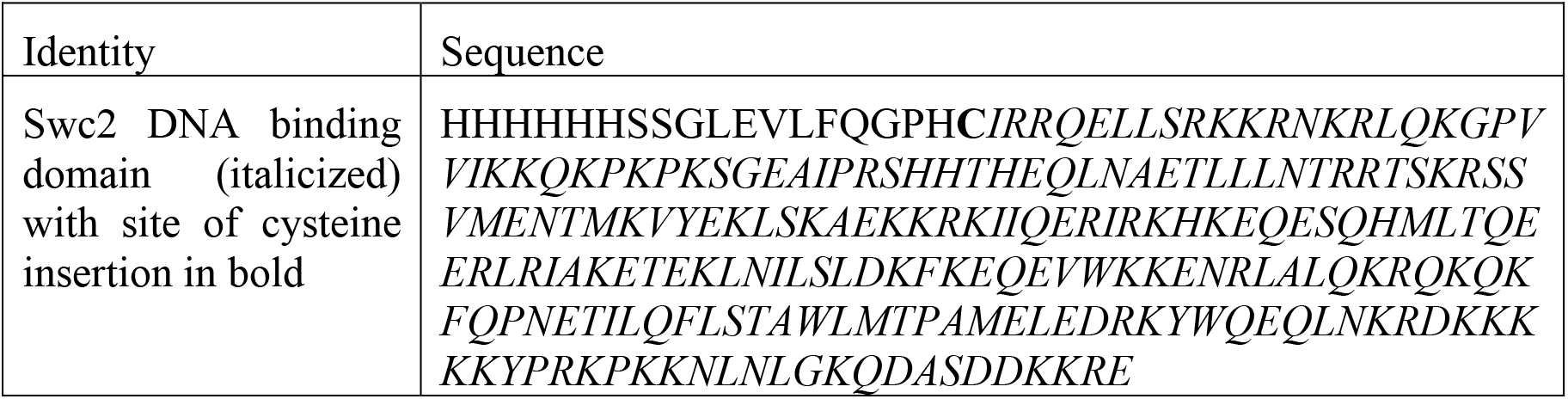
Protein construct sequence.

### dCas9 crRNAs, fluorescent tracrRNA annealing, and RNP assembly

dCas9 was purchased from Integrated DNA Technologies (IDT), as Alt-R S.p.d Cas9 Protein V3 and stored at -80°C until Ribonucleoprotein (RNP) assembly. crRNAs used to target 5 sites along lambda DNA were ordered from IDT. The crRNAs used were previously validated (Sternberg *et al*., 2014) and are listed in **Table 2**. Custom 3’-amine modified tracrRNA was ordered from IDT and reacted with mono-reactive NHS-ester Cy5 dye [Fisher Scientific cat# 45-001-190]. The labeled product was reverse-phase HPLC purified. crRNA and Cy5-tracrRNA was annealed in IDT duplex buffer (cat# 11-01-03-01) in equimolar amounts by heating the mixture to 95°C for 5 minutes and allowing it to cool to room temperature slowly on the benchtop. RNP complexes were assembled by mixing annealed guide RNA and dCas9 in a 1.5:1 molar ratio and allowing the mixture to stand at room temperature for 15 minutes prior to use. Aliquoted RNPs were flash frozen and stored at -80°C until time of use. Buffers for RNP assembly and cryo-storage are the same and contains: 20 mM Tris-HCl pH 7.5, 200 mM KCl, 5% glycerol, and 1 mM TCEP. dCas9 RNPs were diluted to 10 nM just prior to imaging in 1x NEB 3.1 (cat# B7203S).

**Table 2.** crRNA sequences for dCas9 binding and custom oligos sequences for DNA tethering.

| Identity | Sequence |
| --- | --- |
| Cas9 crRNA sequence “lambda 1” | 5'- /AltR 1 /rGrGrC rGrCrA rUrArA rArGrA<br>rUrGrA rGrArC rGrCrG rUrUrU rUrArG rArGrC<br>rUrArU rGrCrU / AltR2/ -3' |
| Cas9 crRNA sequence “lambda 2” | 5'- / AltR 1 /rGrUrG rArUrA rArGrU rGrGrA<br>rArUrG rCrCrA rUrGrG rUrUrU rUrArG rArGrC<br>rUrArU rGrCrU / AltR2/ -3' |
| Cas9 crRNA sequence “lambda 3” | 5'- / AltR 1 /rCrUrG rGrUrG rArArC rUrUrC<br>rCrGrA rUrArG rUrGrG rUrUrU rUrArG rArGrC<br>rUrArU rGrCrU / AltR2/ -3' |
| Cas9 crRNA sequence “lambda 4” | 5'- /AltR1 /rCrArG rArUrA rUrArG rCrCrU<br>rGrGrU rGrGrU rUrCrG rUrUrU rUrArG rArGrC<br>rUrArU rGrCrU / AltR2/ -3' |
| Cas9 crRNA sequence “lambda 5” | 5'- /AltR 1 /rGrGrC rArArU rGrCrC rGrArU<br>rGrGrC rGrArU rArGrG rUrUrU rUrArG rArGrC<br>rUrArU rGrCrU / AltR2/ -3' |
| 3x-biotin-cos1 oligo | 5' - /5Phos/ AGG TCG CCG CCC<br>TT/iBiodT/TT/iBiodT/TT/3BiodT/-3' |
| 3x-biotin-cos2 oligo | 5'- /5Phos/ GGG CGG CGA CCT<br>TT/iDigN/TT/iDigN/TT/3DigN/-3' |

### Lambda DNA preparation

Biotinylated lambda DNA used in SWR1 sliding on naked DNA assays was purchased from LUMICKS (SKU: 00001). Lambda DNA used in nucleosome array assays was made with 3 biotins on one end, and 3 digoxigenin on the other end using the following protocol. Custom oligos were ordered from IDT with sequences listed in **Table 2**. Lambda DNA was ordered from NEB (cat# N3011S). Oligo 1 was annealed to lambda DNA by adding a 25-fold molar excess of oligo to lambda DNA, in an annealing buffer containing 30 mM HEPES pH 7.5 and 100 mM KCl. This mixture was heated to 70°C for 10 minutes and allowed to cool slowly to room temperature on the benchtop. 2 µL of NEB T4 DNA ligase (400U, cat# M0202S) was added along with T4 DNA ligase buffer containing ATP and allowed to incubate at room temperature for 30 minutes. Then 50-fold molar excess of oligo 2 was added to the mixture along with an additional 1 µL of T4 DNA ligase and T4 DNA ligase buffer (NEB) with ATP adjusting for the change in volume and allowed to incubate at room temperature for 30 minutes. The resulting mixture was heat inactivated at 65°C for 10 minutes. End-labeled lambda DNA was purified using Qiaex II gel-extraction DNA clean-up kit following the manufactures’ instructions (Qiagen cat# 20021).

### Lambda nucleosome array construction and validation

A salt gradient dialysis approach was used to reconstitute nucleosomes onto lambda DNA using methods optimized in the lab based on previously established protocols (Luger et al., 1999; Vary et al., 2003). Buffers used in this reconstitution are as follows: high salt buffer [10 mM Tris-HCl pH 7.5, 1 mM EDTA pH 8, 2 M NaCl, 0.02% NP-40, 5 mM 2-Mercaptoethanol (BME)], and low salt buffer [10 mM Tris-HCl pH 7.5, 1 mM EDTA pH 8, 50 mM NaCl, 0.02% NP-40, 5 mM BME]. Cy5-labeled H3 containing octamer, with the same composition and preparation as previously used (Ranjan *et al*., 2013), was titrated onto the lambda DNA in the follow molar ratio to DNA: [10:1, 50:1, 100:1, 200:1, 500:1, 700:1]. Reconstitution reactions were prepared in 10 mM Tris pH 7.5, 1 mM EDTA pH 8, 0.1 mg/mL BSA Roche (cat # 10711454001), 5 mM BME. Any dilutions of octamer were prepared in octamer refolding buffer: [10 mM Tris-HCl pH 7.5, 1 mM EDTA pH 8, 2 M NaCl, 5 mM 2-Mercaptoethanol (BME)]. A 16-hour dialysis was set-up by placing the reconstitution mixture in a 7 kDa MWCO Slide-A-Lyzer MINI Dialysis Device (Thermo Scientific cat # 69560) and placed in a flotation device in high-salt buffer. Low-salt buffer was slowly dripped into high-salt buffer for the duration of the dialysis with constant stirring. At the end of this dialysis period, the dialysis solution was dumped and replaced by 100% low-salt buffer and allowed to dialyze for an additional hour. The reconstitution efficiency was first assessed using an electrophoretic mobility shift assay (EMSA) (**Figure 6–figure supplement 1**). Lambda nucleosome arrays were loaded on a 0.5% agarose gel made with Invitrogen UltraPure Agarose (fisher scientific cat # 16-500-500) and 0.25x TBE. Sucrose loading buffer without added dyes was used to load samples on the gel. The gel was run for 1 hour and 45 minutes at 100V in 0.25x TBE.

Arrays contained a variable number of nucleosomes, where the mean number of nucleosomes per array is 40 ± 5 (standard deviation) for a total of 19 arrays. The number of nucleosomes per array was estimated from the length of the lambda nucleosome array at 5 pN force before and after nucleosome unwrapping. On average, approximately 34.6 nm of lengthening at 5pN corresponded to the unwrapping of a single nucleosome, therefore the difference in length before and after unwrapping was used to estimate the number of nucleosomes per array.

### Dual optical tweezers and confocal microscope set-up and experimental workflow

The LUMICKS cTrap (series G2) was used for optical tweezer experiments, configured with two optical traps. The confocal imaging laser lines used were 532 nm (green) and 640 nm (red) in combination with emission bandpass filters 545-620 nm (green) and 650-750 nm (red). A C1 type LUMICKS microfluidics chip was used. The microfluidics system was passivated at the start of each day of imaging as follows: 0.1% BSA was flowed at 0.4 bar pressure for 30 minutes, followed by a 10-minute rinse with PBS at 0.4 bar pressure, followed by 0.5% Pluronic F-127 flowed at 0.4 bar pressure for 30-minutes, followed by 30-minute rinse with PBS at 0.4 bar pressure. For SWR1 sliding on naked DNA, 4.2 µm polystyrene beads coated in streptavidin (Spherotech cat# SVP-40-5) were caught in each trap, and LUMICKS biotinylated lambda DNA was tethered. Both traps had trap stiffness of about 0.8 pN/nm. For SWR1 sliding on lambda nucleosome array, a 4.2 µm polystyrene bead coated in streptavidin was caught in trap 1, and a 2.12 µm polystyrene bead coated in anti-digoxigenin antibody (Spherotech cat# DIGP-20-2) was caught in trap 2 which is upstream in the path of buffer flow to trap 1. For this configuration, trap 1 had a trap stiffness of about 0.3 pN/nm whereas trap 2 had a trap stiffness of about 1.2 pN/nm. The presence of a single tether was confirmed by fitting a force extension plot to a worm like chain model in real time while collecting data using LUMICKS BlueLake software. For confocal scanning, 1.8 µW of green and red laser power were used. For most traces, the frame rate for SWR1 imaging was 50 msec, whereas for Swc2 it was 20 msec. Experiments were performed at room temperature. SWR1 and Swc2 were both imaged in histone exchange reaction buffer [25 mM HEPES pH 7.6, 0.37 mM EDTA, 5% glycerol, 0.017% NP40, 70 mM KCl, 3.6 mM MgCl_2_, 0.1 mg/mL BSA, 1 mM BME] made in imaging buffer. dCas9 was added to the flow chamber in Cas9 binding buffer [20 mM Tris-HCl pH 8, 100 mM KCl, 5 mM MgCl_2_, 5% glycerol] made in imaging buffer. Imaging buffer [saturated Trolox (Millipore Sigma cat# 238813), 0.4% dextrose] is used in place of water when preparing buffers. All buffers were filter sterilized with a 0.2 μm filter prior to use.

### TIRF based binding kinetics assay and analysis

We co-localized SWR1 binding to Cy5-labeled dsDNAs of different lengths for real-time binding kinetic measurements (**Figure 1–figure supplement 1D-E**). These experiments were all conducted using flow cells made with PEG-passivated quartz slides using previously detailed methods (Roy et al., 2008). The appropriate biotinylated Cy5-labeled DNA was immobilized on the surface of the PEG-passivated quartz slide using neutravidin. After DNA immobilization, the channels of the flow cell were washed to remove free DNA and imaging buffer was flowed into the channel. Next, 5 nM Cy5-SWR1 in imaging buffer was flowed into the channel immediately after starting image acquisition. A standard smFRET imaging buffer with oxygen scavenging system was used as has been previously established (Joo and Ha, 2012). The first 10 frames (1s) of each imaging experiment were collected using Cy5-excitation so that all Cy5-DNA spots could be identified. The remaining 299 seconds of the movie were collected under Cy3-excitation so that Cy3-SWR1 could be imaged. Data analysis was carried out using homemade IDL scripts for image analysis and MATLAB scripts for data analysis. The data was analyzed so that all the Cy5-DNA molecules in an image were identified from the first second of the movie under Cy5-excitation. Next, the Cy3 intensity was monitored for the remainder of the movie for each DNA molecule. SWR1 binding to nucleosomes was detected by a sharp increase in Cy3 signal in spots that had Cy5 signal.

The on-rate was defined as the time between when Cy3-SWR1 was injected into the imaging chamber to when Cy3-SWR1 first bound to a specific DNA molecule resulting in an increase in Cy3 intensity. The off-rate was defined as the length of time Cy3-SWR1 was bound to a DNA molecule which is the duration of the high Cy3 fluorescence state. While only one on-rate measurement could be conducted for one DNA molecule, multiple off-rate measurements could be made as one DNA molecule was subjected to multiple Cy3-SWR1 binding events. Binding events where more than one SWR1 were bound to the DNA were excluded from the off-rate analysis. Off-rate measurements under different laser intensities were made by measuring the laser power immediately prior to the imaging experiment (**Figure 1–figure supplement 1C**). All experiments were conducted using imaging channels from the same quartz slide to minimize differences in laser intensity that can result from changes in shape of the TIRF spot.

### Single particle tracking and data analysis

LUMICKS Bluelake HDF5 data files were initially processed using the commercial Pylake Python package to extract kymograph pixel intensities along with corresponding metadata. Particle tracking was then performed in MATLAB (MathWorks). First, spatially well-separated particles were individually segmented from full-length kymographs containing multiple diffusing particles. Next, for each time-step, a one-dimensional gaussian was fit to the pixel intensities to extract the centroid position of the particle in time. Then the MSD for each time-lag was calculated using:

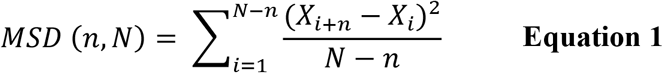

where N is the total number of frames in the trace, n is the size of the time lag over which the MSD is calculated, i is the sliding widow over which displacement is measured, X is the position of the particle. Since particles exhibit Brownian diffusion, the diffusion coefficient for each particle was then calculated from a linear fit to the initial portion of the mean squared displacement (MSD) versus time lag plot by solving for D using: *MSD* = 2*DT*. For mean MSD plots, traces with the same frame rate were averaged together, resulting in a slightly different n-value as compared to all trajectories in a condition.

For the linear fit, the number of points included varied to optimize for a maximal number of points fit with the highest Pearson correlation (r^2^) and a p-value lower than 0.05. For particles where this initial best fit could not be found, the first 25% of the trace was linearly fit. Fits that produced negative slope values corresponded to traces where particles are immobile; to reflect this, negative slopes were given a slope of 0. Finally, outlier traces with diffusion coefficients greater than 0.14 µm^2^/s for SWR1 or 5 µm2/s for Swc2 were dropped; in every case this consisted of less than 3% of all traces. The distribution of diffusion coefficients estimated using this method was almost identical to what is produced using an alternative method which extracts diffusion coefficients using a linear fit from time lags 3-10 rejecting fits with r^2^< 0.9 (Tafvizi *et al*., 2008) (data not shown). A summary of statistics as well as criteria for excluding traces is provided in **Table 3**. Also included are the number of biological and technical replicates per condition. A biological replicate is defined as a fresh aliquot of protein imaged on a different imaging day, whereas a technical replicate is the number of distinct DNAs or nucleosome arrays used per imaging condition; a single DNA could accommodate one or more fluorescently tagged proteins.

**Table 3.**
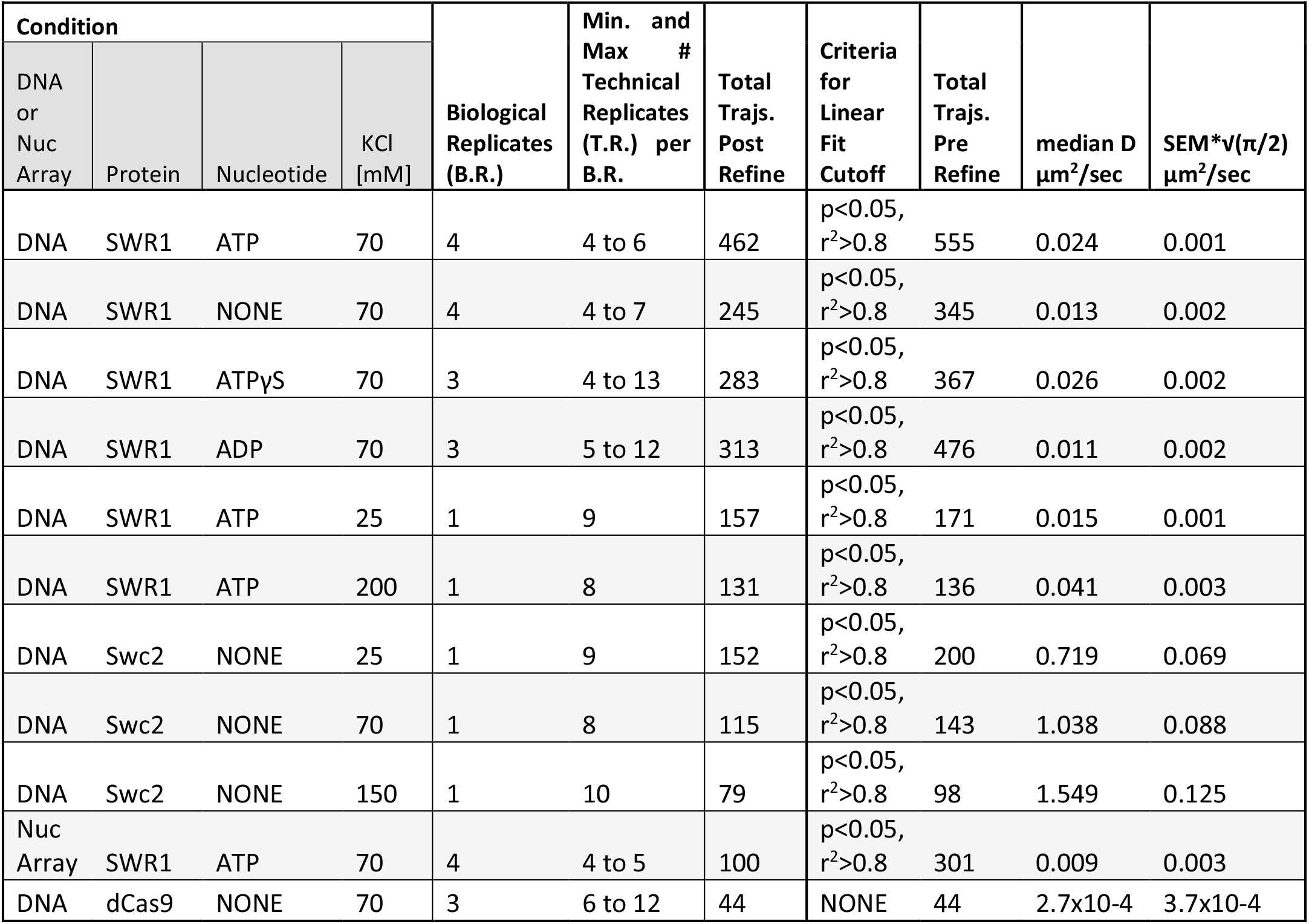
Summary of median diffusion coefficients as well as rejection criteria implemented per condition for particle refinement. Also included is information regarding biological and technical replicates. ‘Trajs.’ stands for trajectories.

| Condition | | | | Biological Replicates (B.R.) | Min. and Max # Technical Replicates (T.R.) per B.R. | Total Trajs. Post Refine | Criteria for Linear Fit Cutoff | Total Trajs. Pre Refine | median D $\mu\text{m}^2/\text{sec}$ | SEM*$\sqrt{(\pi/2)}$ $\mu\text{m}^2/\text{sec}$ |
| --- | --- | --- | --- | --- | --- | --- | --- | --- | --- | --- |
| DNA or Nuc Array | Protein | Nucleotide | KCl [mM] |  |  |  |  |  |  |  |
| DNA | SWR1 | ATP | 70 | 4 | 4 to 6 | 462 | $p < 0.05$, $r^2 > 0.8$ | 555 | 0.024 | 0.001 |
| DNA | SWR1 | NONE | 70 | 4 | 4 to 7 | 245 | $p < 0.05$, $r^2 > 0.8$ | 345 | 0.013 | 0.002 |
| DNA | SWR1 | ATPyS | 70 | 3 | 4 to 13 | 283 | $p < 0.05$, $r^2 > 0.8$ | 367 | 0.026 | 0.002 |
| DNA | SWR1 | ADP | 70 | 3 | 5 to 12 | 313 | $p < 0.05$, $r^2 > 0.8$ | 476 | 0.011 | 0.002 |
| DNA | SWR1 | ATP | 25 | 1 | 9 | 157 | $p < 0.05$, $r^2 > 0.8$ | 171 | 0.015 | 0.001 |
| DNA | SWR1 | ATP | 200 | 1 | 8 | 131 | $p < 0.05$, $r^2 > 0.8$ | 136 | 0.041 | 0.003 |
| DNA | Swc2 | NONE | 25 | 1 | 9 | 152 | $p < 0.05$, $r^2 > 0.8$ | 200 | 0.719 | 0.069 |
| DNA | Swc2 | NONE | 70 | 1 | 8 | 115 | $p < 0.05$, $r^2 > 0.8$ | 143 | 1.038 | 0.088 |
| DNA | Swc2 | NONE | 150 | 1 | 10 | 79 | $p < 0.05$, $r^2 > 0.8$ | 98 | 1.549 | 0.125 |
| Nuc Array | SWR1 | ATP | 70 | 4 | 4 to 5 | 100 | $p < 0.05$, $r^2 > 0.8$ | 301 | 0.009 | 0.003 |
| DNA | dCas9 | NONE | 70 | 3 | 6 to 12 | 44 | NONE | 44 | $2.7 \times 10^{-4}$ | $3.7 \times 10^{-4}$ |

We estimated the localization precision using the following formula:

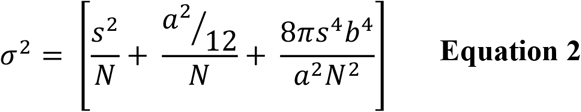

where N is the number of photons collected which was on average 12.9 photons per 5-pixel window surrounding the centroid (data not shown); s is the standard deviation of the microscope point-spread function, 294 nm; a is the pixel size, 100 nm; and b is the background intensity which was on average 0.8 photons per 5-pixel window. This results in a σ = 82 nm.

### Calculation of theoretical maximal hydrodynamic diffusion coefficients

The radius of gyration of SWR1 and Swc2 were calculated using the following formulas. First, the volume (V) of each particle was estimated using the following equation:

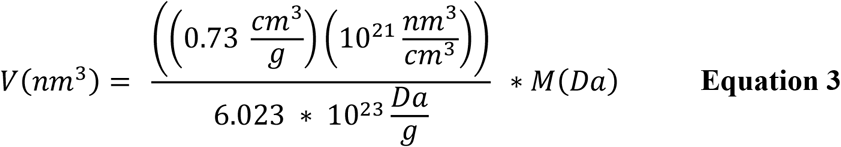

Then, the radius of gyration was estimated using the following equation:

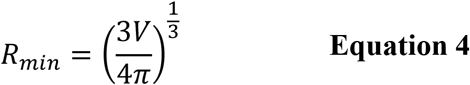

where M is mass in Daltons (Erickson, 2009). Given the input of 1 MDa for SWR1 and 25.4 kDa for Swc2, the resulting radii of gyration are 6.62 nm SWR1 and 1.94 nm for Swc2. Next, the theoretical upper limit of 1D diffusion with no rotation was calculated using the following formula:

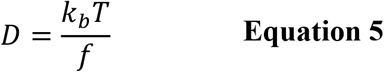

Where:

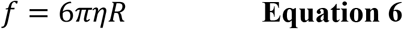

and η is the viscosity 9×10^−10^ pN*s/nm^2^ (Schurr, 1979). The resulting upper limit without rotation for SWR1, is 36.7 µm^2^/s and for Swc2 it is 125 µm^2^/s. When computing the upper limit of 1D diffusion with rotation, the following formula considers the energy dissipation that comes from rotating while diffusing:

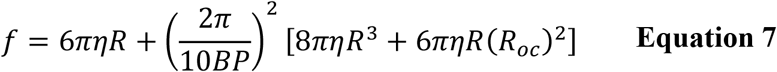

where R_oc_ is the distance between the center of mass of the DNA and the bound protein, and 10 BP is the length of one helical turn or 3.4 nm (Ahmadi *et al*., 2018; Bagchi et al., 2008; Blainey *et al*., 2009). Since we do not have structures of SWR1 or Swc2 bound to dsDNA alone, we report both the maximal and minimal value of the theoretical upper limit, where the minimal value corresponds to R_oc_ = R and the maximal value corresponds to R_oc_ = 0. For SWR1 this minimum value is 0.105 µm^2^/s and the maximum value is 0.183 µm^2^/s whereas for Swc2 this minimum value is 4.01 µm^2^/s and the maximum value is 6.86 µm^2^/s.

### Scanning speed estimation

Lambda DNA tethered at its ends to two optically trapped beads was pulled to a tension of 5 pN, which resulted in a length approximately 92% of its contour length (15.2 µm). The length per base pair of DNA, 0.31 nm, is therefore slightly shorter than the value at full contour length (Baumann *et al*., 2000). The length of the NDR, 150 bp, in our conditions is therefore roughly 0.047 µm long. Since our localization precision is low, ∼82 nm (**see Equation 2**), we do not have diffusion information at the resolution of base pairs, and therefore do not consider discrete models to approximate scanning speed. Given a median diffusion coefficient of SWR1 in the presence of 1 mM ATP of 0.024 µm^2^/sec, and the one-dimensional translational diffusion, *l*= 2*DT*, where *l*is the length in µm of DNA, we can approximate the time required to scan this length of DNA to be 0.093 seconds assuming a continuous model (Berg, 1983).

## Author contributions

C.C.C., M.F.P. C.W. and T.H. conceived the project. C.C.C. designed, performed, and analyzed cTrap experiments, M.F.P. designed, performed, and analyzed TIRF experiments. G.P. and R.K.L. purified site specifically labeled SWR1, M.F.P. purified Swc2. T.Z. assisted with cTrap measurements. C.C.C. wrote the manuscript with contributions from all authors. C.W. and T.H. supervised the project.

## Acknowledgements

We thank Dr. Kelsey Bettridge for providing a template script for single particle tracking in Matlab.

## Data and materials availability

Raw data has been uploaded to Dryad in the form of a Matlab structured arrays. All Matlab codes used to generate the main figures are publicly available at https://github.com/ccarcam1/SWR1_1D_Diffusion_Publication

