## Supplementary Information for "ATP Binding Facilitates Target Search of SWR1 Chromatin Remodeler by Promoting One-Dimensional Diffusion on DNA"

**Figure 1–figure supplement 1**

**Figure 2–figure supplement 1**

**Figure 4–figure supplement 1**

**Figure 4–figure supplement 2**

**Figure 5–figure supplement 1**

**Figure 6–figure supplement 1**

**Figure 6–figure supplement 2**

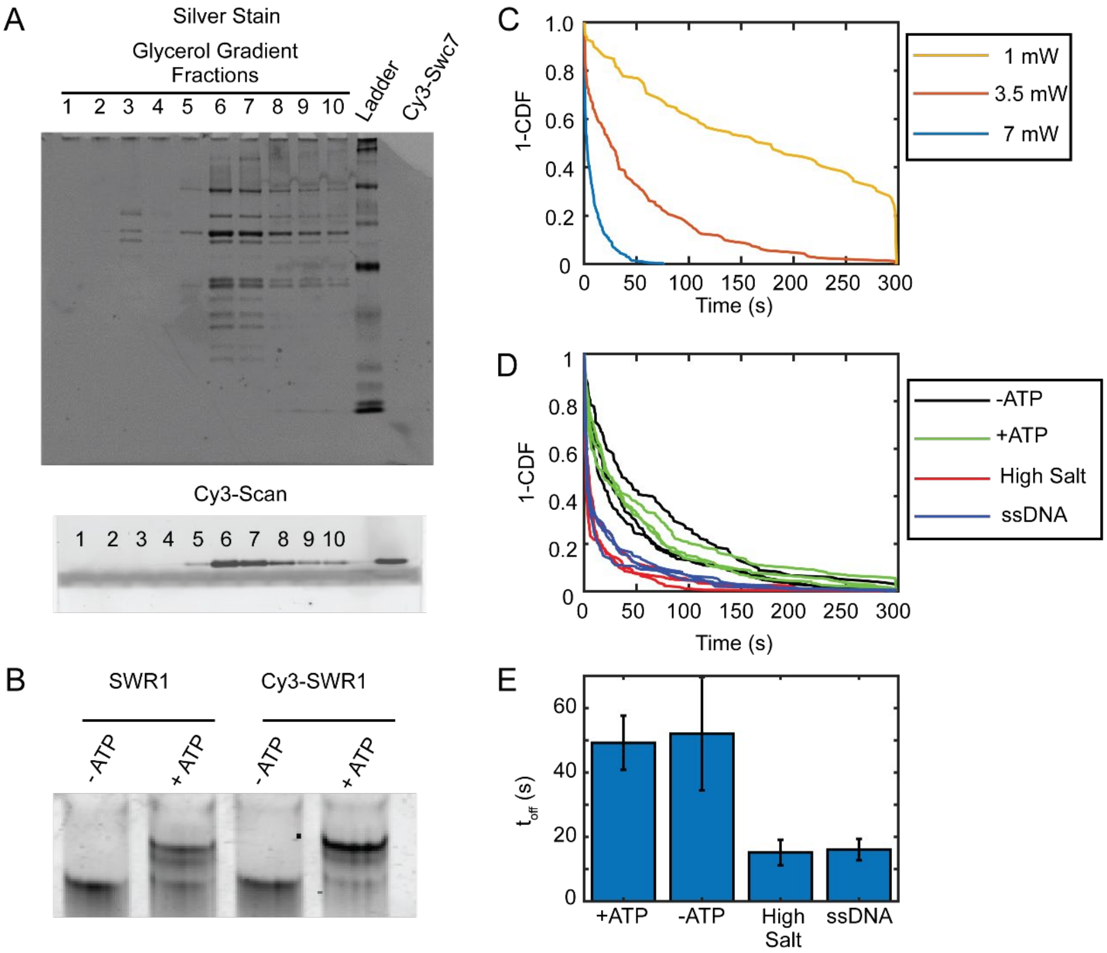


**Figure 1–figure supplement 1. Cy3-SWR1 purification and DNA binding kinetics.**
(A) A glycerol gradient purification of the Cy3-SWR1 complex after Cy3-Swc7 reincorporation. A silver stain (top image) shows that the SWR1 complex eluted in fractions 6 and 7 of the gradient. A Cy3 image of the same gel shows that Cy3-Swc7 also is found in fractions 6 and 7 (confirmed by a Cy3-Swc7 only control at the end of the gel). This demonstrates that the Cy3-Swc7 is incorporated into the SWR1ΔSwc7 complex. (B) A histone exchange assay that shows Cy3-SWR1 (right lanes) is as active as the wild type SWR1 complex. The gel-shift is caused by the incorporation of triple flag-tagged ZB dimers into the nucleosome. (C) $t_{off}$ for Cy3-SWR1 bound to 150 bp DNA measured at different laser powers. These measurements show that the lifetime of Cy3-SWR1 bound to 150 bp DNA is photobleaching limited. (n = 1 measurement for each condition) (D) 1-CDF for Cy3-SWR1 bound to 150 bp DNA and exposed to ATP, 200 mM NaCl and 0.2 ug/mL of salmon sperm DNA (ssDNA). Three technical replicates are shown. (n = 3 measurements for each condition, all data is shown) (E) $t_{off}$ for Cy3-SWR1 bound to 150 bp DNA in different conditions, same as those shown in panel (D) (n = 3, error bars represent standard deviation).

**Figure 1–figure supplement 1– source data 1**
Gel images (Coomassie and Cy3 scans) shown in panels A and B

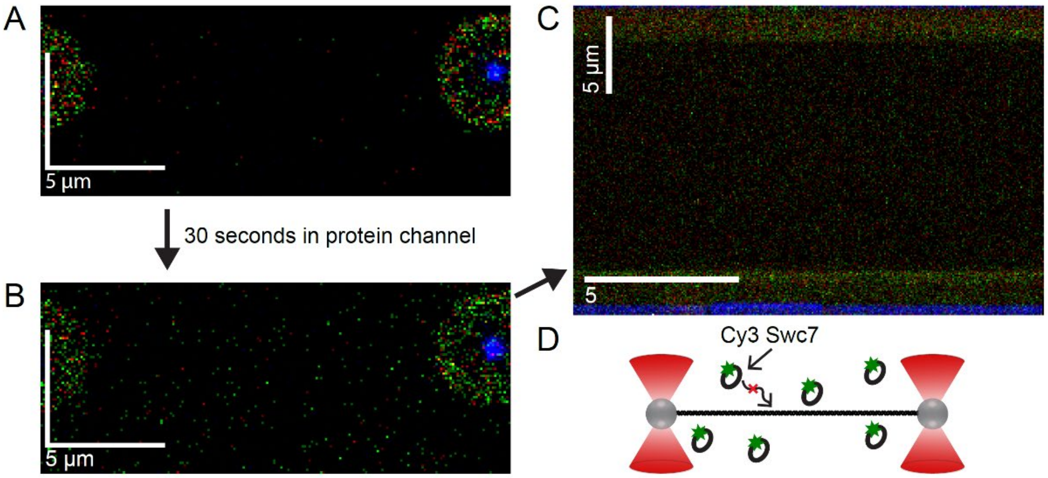


**Figure 2–figure supplement 1. Swc7 cy3 does not bind DNA without the SWR1 complex.** (A) A scan of lambda DNA pulled to 5pN of tension in the absence of Cy3 labeled Swc7 imaged with green excitation. (B) A scan of lambda DNA in Cy3 Swc7 protein channel. Protein solution was flowed briefly to refresh the protein solution in the channel before and while DNA is introduced into the channel. The amount of Cy3 labeled Swc7 is equimolar to what is used for SWR1 sliding experiments. Apart from increased background, there is no specific interaction of Cy3 Swc7 with the DNA alone. (C) A kymograph along the length of the lambda DNA at a time resolution of 0.05 sec per line scan also shows no bound Cy3 Swc7. (D) Schematic summary: Cy3 Swc7 doesn't bind lambda DNA alone, even at the faster time resolution used to generate a kymograph.

**Figure 2–figure supplement 1– source data 1**

Raw scans and kymograph tiff files

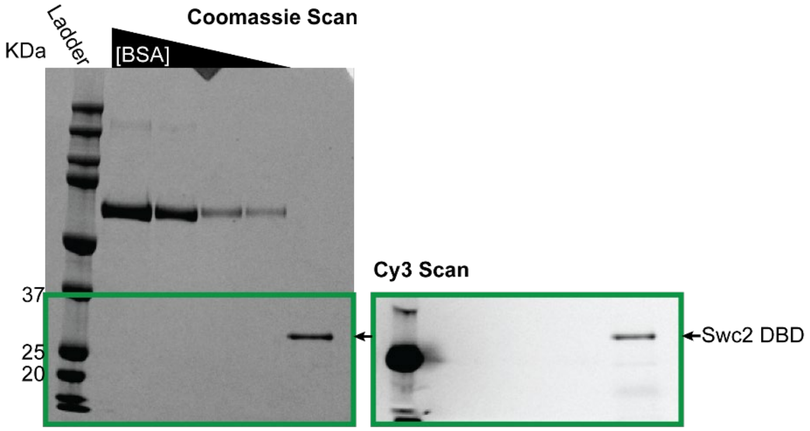


**Figure 4–figure supplement 1. Purification and fluorophore labeling of the Swc2 DBD.** Denaturing SDS PAGE gel of the Swc2 DNA binding domain (DBD) after nickel his-tag purification and labeling with Cy3-maleimide. A Cy3 scan of the gel (right) reveals that the Swc2 DBD (single band in Coomassie stain on the left) has been labeled with Cy3.

**Figure 4–figure supplement 1– source data 1**

Gel images (Coomassie and Cy3 scans)

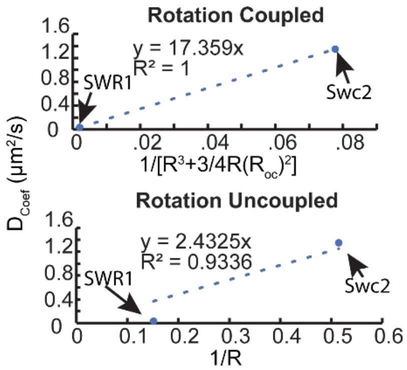


**Figure 4–figure supplement 2. Rotation coupled vs uncoupled diffusion models and protein size effects on diffusion coefficient.** Apparent diffusion coefficient as a function of two relationships to the radius of SWR1 or Swc2 DBD where the protein size is printed below the graph and the corresponding model is stated above. A line passing through the origin is fit to both relationships, and the Pearson correlation coefficient is printed within the graph showing a slightly better fit of the measured diffusion coefficients to the rotation coupled diffusion model.

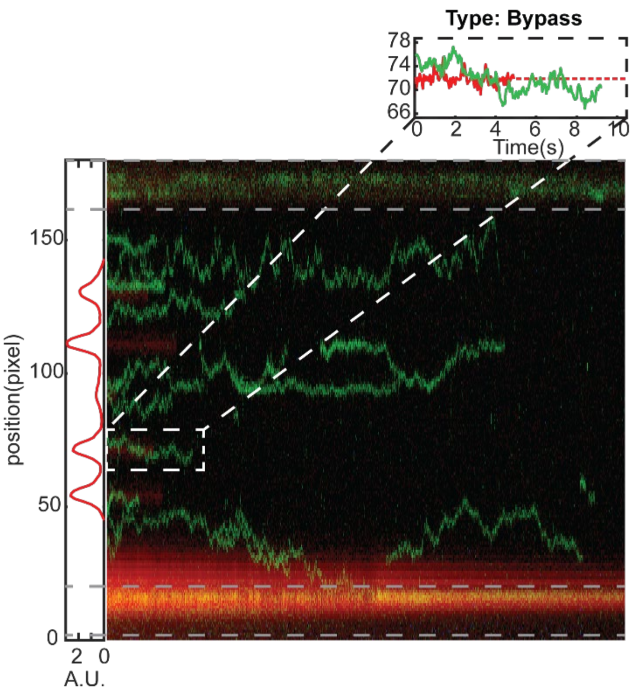


**Figure 5–figure supplement 1. SWR1 bypassing dCas9 was a rare event.** Of 106 total colocalization events, only 3 displayed some form of bypass event. Left of the representative kymograph is a sum of the red intensity, which is used to calculate the centroid of dCas9 for colocalization analysis.

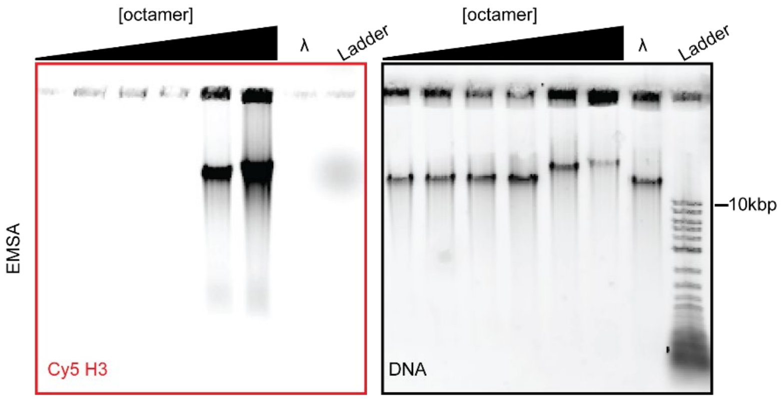


**Figure 6–figure supplement 1. Lambda nucleosome array EMSA.** Cy5 labeled H3 octamer (~20% labeling efficiency) was used in the reconstitution of nucleosomes onto lambda DNA via salt gradient dialysis. Typhoon imager scans (left Cy5 scan, right SYBR Gold scan). Octamer concentrations used are as follows reported as molar ratio of octamer to DNA, from left to right: 10:1, 50:1, 100:1, 200:1, 500:1, 700:1. Lambda DNA alone is shown for reference. The 700:1 condition was selected for use in experiments.

**Figure 6–figure supplement 1– source data 1**

Gel images (Coomassie and Cy3 scans)

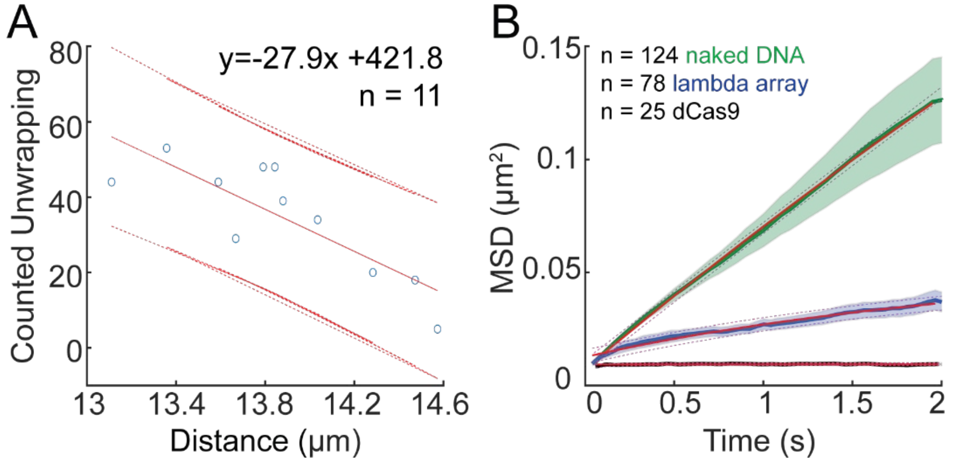


**Figure 6–figure supplement 2. Lambda nucleosome array nucleosome quantification and separation estimation.** (A) The relationship between the length of the nucleosome array at 5pN and the number of unwrapping events counted. (B) Mean MSD vs time is fit to y = m*(1-exp(-b*x))+c. For SWR1 on the lambda nucleosome array [blue], the limit approached 0.054 $\mu m^2$, whereas SWR1 diffusing on lambda DNA [green] produced a limit 1.1 $\mu m^2$, and dCas9 [black] produced a limit of 0.009 $\mu m^2$ which is within the limit of resolution. Fits and 95% confidence intervals are overlayed.

**Figure 6–figure supplement 2– source data 1**

Data underlying panel A
